# Time-Resolved Neuronal Network Dynamics Distinguish Pathological States in Organoid Models

**DOI:** 10.64898/2026.03.25.714227

**Authors:** Colin M. McCrimmon, Prateik Sinha, Qing Cao, Tonmoy Monsoor, Kartik Sharma, Mehmet Yigit Turali, Ranmal A. Samarasinghe, Vwani Roychowdhury

**Author notes:** Indicates equal contribution (Colin M. McCrimmon, Prateik Sinha, Qing Cao, Tonmoy Monsoor). This work was funded by the Broad Stem Cell Research Center (BSCRC) Innovation Award (R.A.S.) and Transformative Technology Development (TTD) Award (R.A.S. and V.R.), as well as The Michael R. Bloomberg Revocable Trust (R.A.S.).

## Abstract

Human brain assembloids offer a powerful platform for modeling neurological diseases, yet comprehensive methods for analyzing their complex network dynamics are lacking. Here, we developed a time-resolved network analysis pipeline that extracts quantitative biomarkers from two-photon calcium imaging, enabling the detection of subtle differences between disease and control models. We applied this pipeline to assembloids containing a pathogenic *MAPT* p.R406W variant—clinically associated with an Alzheimer’s disease-like phenotype—and their isogenic controls. Our analysis revealed that *mutant* networks exhibit significantly increased degree variance and clustering. This indicates a “hub-like”, interconnected topology prone to hypersynchrony, a finding that parallels the network hyperexcitability and seizure-like features observed in *in-vivo* models of Alzheimer’s disease. Furthermore, a Random Forest classifier trained on these dynamic network features distinguished between diseased and control states with high accuracy (F1 score = 0.90). These results establish that dynamic network properties can serve as potent biomarkers for identifying pathological states in assembloid models, providing a quantitative framework to investigate disease mechanisms and potential therapeutic interventions.

## 1. INTRODUCTION

Human brain organoids, and their fused “assembloid” counterparts, provide a revolutionary platform for studying neurological disease. By recapitulating human-specific aspects of neurodevelopment—including layered cytoarchitecture, diverse cell populations, and complex neuronal activity [1, 2, 3, 4]—these reproducible and scalable 3D models overcome many limitations of traditional animal systems for investigating the functional properties of human neuronal networks. While prior work has demonstrated that these models exhibit rich, spontaneous network activity, the field has lacked standardized, comprehensive, and high-throughput analysis methods needed to fully leverage this technology. This analytical gap is particularly challenging because a primary application of organoids is modeling genetic diseases [5, 6], often by direct comparison to healthy, isogenic controls [3, 4]. To address this critical gap, we developed a comprehensive, time-resolved network analysis pipeline that defines quantitative functional biomarkers of network activity. Our objective was not only to develop a classifier that could distinguish between disease and control states, but also to identify the salient features—the specific quantitative biomarkers—that drive this distinction. By pinpointing these core features, our approach provides critical insight into the fundamental biological mechanisms of network dysfunction, creating a powerful new tool to guide subsequent cellular-level studies and therapeutic development.

## 2. METHODS: FUNCTIONAL BIOMARKERS FROM ASSEMBLOIDS

### 2.1. Organoids to time-resolved neuronal activity data

#### Assembloid generation

We generated cerebral organoids from two human induced pluripotent stem cell lines: one derived from a patient with a pathogenic *MAPT* p.R406W variant (*mutant*), which produces a clinical phenotype resembling Alzheimer’s Disease, and a matched, CRISPR/Cas9-corrected isogenic control line (*control*) [7]. Using established protocols [3, 4], we directed organoids from each line toward either a hippocampal (Hc) or ganglionic eminence (GE) fate. At 56 days in vitro (DIV), we fused Hc and GE organoids from each line to create *mutant* Hc+GE assembloids and *control* Hc+GE assembloids. We introduced a genetically encoded calcium indicator GCaMP7f at ∼106 DIV via transduction with an AAV-syn-jGCaMP7f virus (Addgene, 104488-AAV1), and then recorded neuronal network activity from 8 *mutant* and 5 *control* assembloids using 2PCI at ∼120 DIV. Ultimately, 175 100-second recordings from these 13 assembloids, which exhibited sufficient neuronal activity for robust network analysis, were included in the subsequent analysis.

To ensure statistical independence of the n=175 recordings, we compared intra-organoid versus inter-organoid variability. Distances between repetitions within a single organoid were statistically indistinguishable from distances across different organoids in our feature space (Kolmogorov-Smirnov test, *p >* 0.3).

#### 2PCI to individual neuron time-series

We extracted activity from regions of interest (i.e. putative neurons) in 2PCI recordings using a custom MATLAB script that leverages the CaImAn toolbox [8]. CaImAn implements a probabilistic decomposition model based on constrained non-negative matrix factorization and Bayesian inference to perform simultaneous source extraction and spike deconvolution. The final processed data consisted of inferred raster plots of action potentials (spikes) from each neuron that were used in subsequent network construction.

### 2.2. Neuronal activity to dynamic correlation networks

Let 𝒳= {*x*_*i*_[*t*] ∈ ℝ | *i* ∈ {1, …, *n*}, *t* ∈ {1, …, *T*}} denote the neuron traces (number of spikes) over *T* = 250 uniformly sampled points, where *i* corresponds to the *i*^*th*^ neuron and *t* corresponds to sample index where one sample corresponds to Δ*t* = 400 ms. We use a sliding window of length *W* = 25 with stride *S* = 1, resulting in 226 windowed data vectors for each neuron. For window *k* starting at *t*_*k*_ = 1 + (*k* − 1)*S*, the windowed data vector for neuron *i* is: 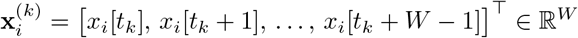.

#### Correlation network for *k*^*th*^ window

We initially construct an undirected, weighted, fully connected network

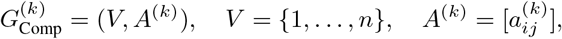

where *A*^(*k*)^ is the weighted adjacency matrix for window *k*. Edge weights, 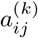, are the sample Pearson correlation coefficients between the *k*^*th*^ windowed data vectors for neurons *i* and 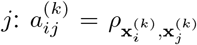. For each *k*, several 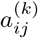 could be negligible, thus we use a percentile-threshold, *τ*, and only retain edges in the top *τ* -percentile. Next, the Giant Connected Component (GCC) of this network is computed and labeled 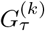. By sweeping across all *k*, we get 226 samples from the underlying network dynamics: {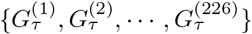}.

### 2.3. Capturing temporal neuronal network dynamics using graph-theoretic measures

#### Network structure descriptors for the *k*^*th*^ window

For network snapshot 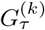 we compute eight local and global descriptors that summarize the network structure (see [9] for definitions of these well-known metrics): (i) normalized size of 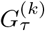; (ii) clustering coefficient; (iii) mean of the node-degree distribution of 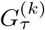; (iv) variance of the node-degree distribution of 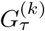; (v) diameter of 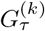; (vi) algebraic connectivity (Fiedler eigenvalue) of 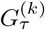; (vii) sum of edge weights of 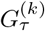; (viii) sum of Euclidean inter-neuron distances within 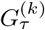, computed from the recorded neuron coordinates.

#### Summarizing temporal network structure descriptors

Let 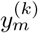 be the *m*-th descriptor measured on 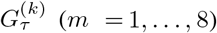 To summarize temporal evolution across *k* = 1, …, *K*, we compute the first four (sample) moments for each scalar graph statistic: mean *µ*_*y*_, variance 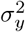, skewness *γ*_*y*_, and kurtosis *κ*_*y*_. Concatenating the four moments across the eight descriptors yields a fixed 32-dimensional signature per assembloid, 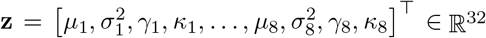, which preserves typical levels and temporal variability or asymmetry of the evolving correlation network and is suitable for downstream classification.

## 3. ASSEMBLOID CLASSIFICATION USING TEMPORAL NEURONAL NETWORK DYNAMICS

Various neurophysiological properties of the assembloid models, including their temporal neuronal network dynamics, may encode disease-state-related information, and could provide critical insights into network-level dysfunction in neurological disorders.

### 3.1. Classification model, performance metrics and feature selection

#### Classification model

We trained a Random Forest (RF) classifier using a nested, stratified 10-fold cross-validation framework. All hyperparameter optimization was performed exclusively on the training data within each fold. To address class imbalance, we applied differential class weighting (2:1 ratio for control:mutant). We opted for stratified 10-fold crossvalidation rather than leave-one-organoid-out validation due to the lack of significant correlation between recordings from the same organoid and the uneven distribution of recordings across the 13 assembloids.

#### Decision rule and performance metrics

From predicted probabilities 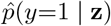 we assign 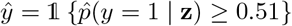 We report accuracy, precision, recall, F1, and ROC–AUC. We also log per-group outcomes (13 organoid groups) to quantify cohort-level consistency.

#### Feature selection

From the initial 32 graph-theoretic statistics, we performed feature selection using importance ranking across folds. The most informative features were the summary statistics of the *GCC node-degree variance* and *transitivity/clustering coefficient*. Group-level comparisons for these features were performed using a two-sided t-test with Bonferroni correction (*α* = 0.05*/*32), remaining significant with *p <* 0.001 (Fig. 3A).

**Fig. 1.**
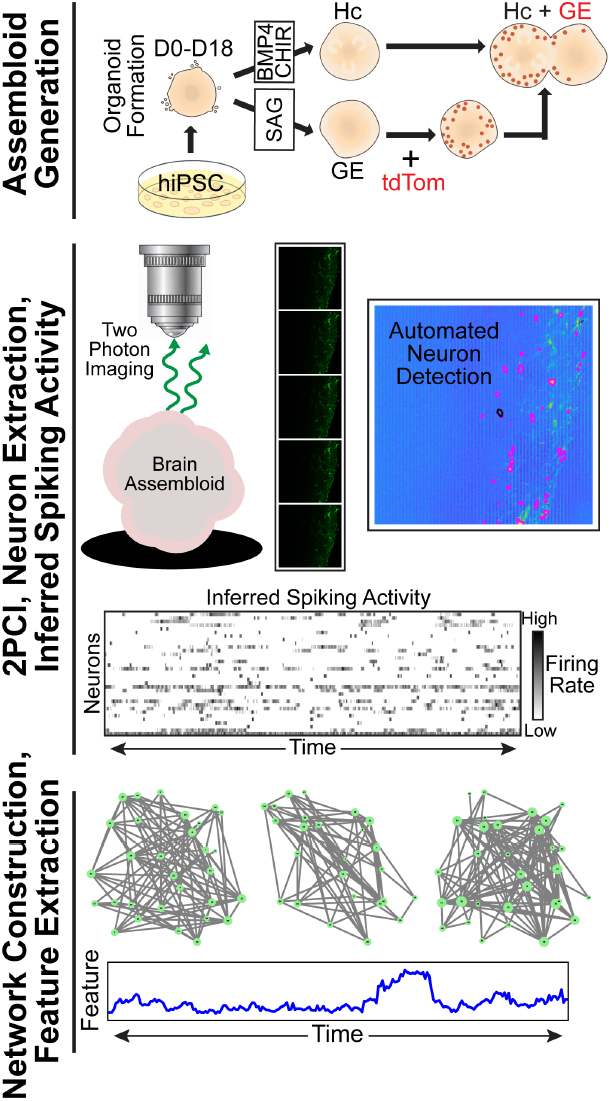
Experimental and analytical workflow. (Top) Organoids were directed towards distinct regional fates starting at 18 DIV and fused into assembloids at 56 DIV. (Middle) Following viral transduction with a genetically encoded calcium indicator, two-photon calcium imaging was performed at 120 DIV. Individual neuronal activity and inferred spike trains were extracted using an automated pipeline. (Bottom) This activity was used to construct time-resolved correlation networks, from which 32 dynamic features were extracted for group-level classification of *mutant* and *control* assembloids.

**Fig. 2.**
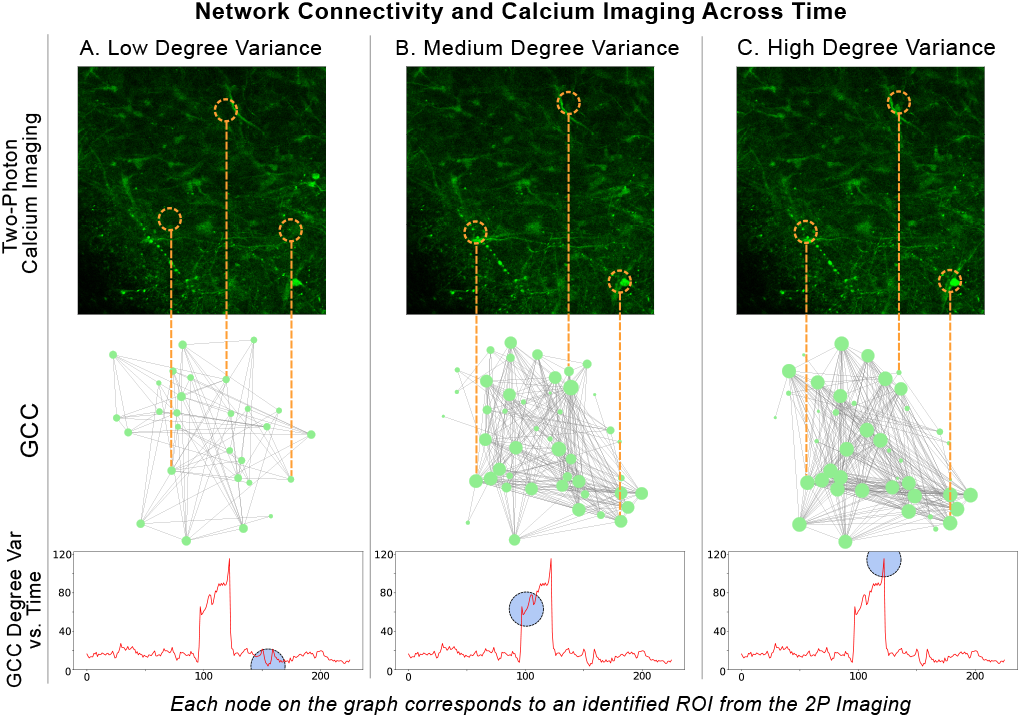
Correlation network connectivity (*τ* = 25) and calcium imaging across time. Each node in the correlation network graph represents a region of interest (putative neuron) extracted from two-photon calcium imaging data. The calcium imaging frame displayed corresponds to the midpoint of the temporal window used to construct the network. Panels A–C depict correlation network states with low, medium, and high GCC degree variance, respectively. The lower panels show the temporal evolution of the GCC degree variance, with the highlighted window (blue) indicating the interval used for graph construction.

**Fig. 3.**
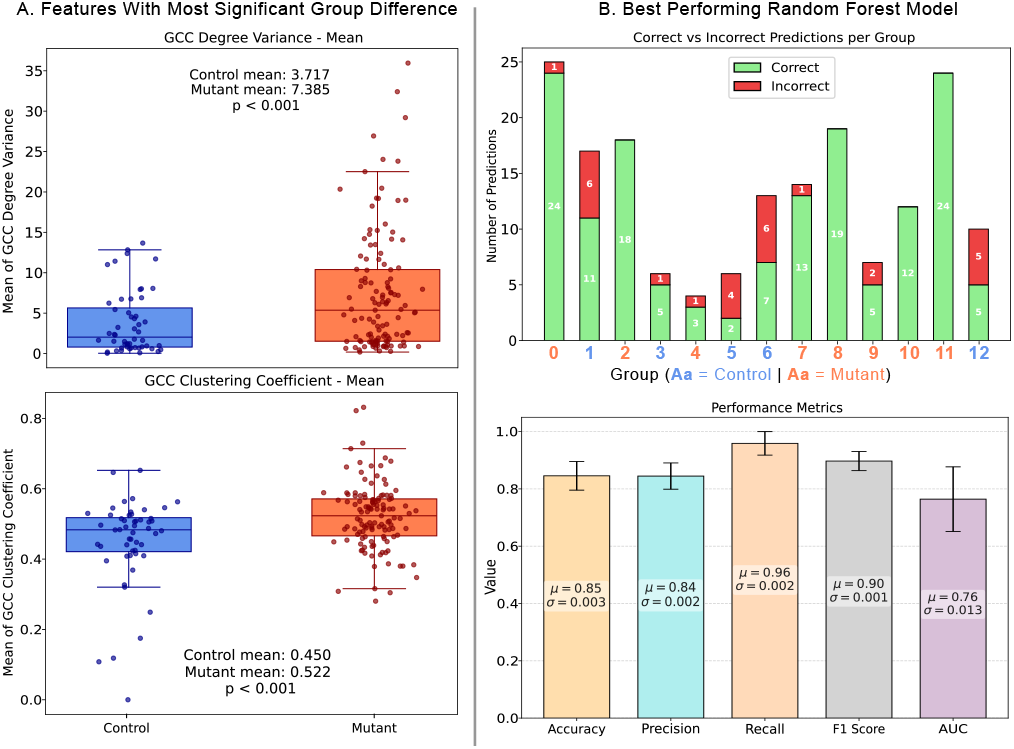
The Random forest (RF) model trained with the summary statistics of GCC node-degree variance and Clustering coefficient performed the best on held-out test set (for details see Section 3.1 on Feature selection for organoid classification). All the plots shown are for the best performing RF model **(A):** Among the four summary statistics of GCC node-degree variance (top panel) and Clustering coefficient (bottom panel), mean had the most significant difference at a group level. **(B)**: (top panel) Model’s predictive accuracy on a per-organoid basis, showing high number of correct classifications (green bars) across most organoids, and few incorrect predictions (red bars) mostly coming from *control* organoids. (bottom panel) Prediction model generalizes well across the 10-folds as shown by the high mean and low variance in the performance metrics distribution across the folds.

### 3.2. Classification results and insights

#### *Mutant* organoids exhibit higher propensity for hypersynchrony: “hub” like structures and higher interconnectedness

Our analysis of network topology revealed that assembloids with the *MAPT* p.R406W variant exhibit significant alterations consistent with a hyperexcitable state. As shown in Fig. 3A, *mutant* networks had a significantly higher mean for node degree variance (top panel), indicating the formation of highly connected “hub” neurons that can drive network-wide synchronization. Concurrently, these *mutant* networks showed a significant increase in the mean clustering coefficient (bottom panel), reflecting a greater degree of local interconnectedness where neighboring neurons are more likely to be connected to each other. These characteristics—the emergence of hubs and tightly clustered local circuits—are hallmarks of a network prone to hypersynchrony. This result is congruent with prior in vivo models of Alzheimer’s Disease [10, 11], where impaired inhibitory interneuron function led to aberrant excitatory activity and epileptiform discharges. The hub-like, highly clustered topology of our *mutant* organoids may therefore be an in vitro manifestation of these imbalanced networks, in which a failure of inhibition drives pathological synchronization.

#### Hypersynchrony dynamics descriptors are sufficient for distinguishing between *mutant* and *control* organoids

The RF predictive model demonstrated strong reliability in classifying organoid state across multiple recordings, correctly identifying the majority of cases for each organoid, as shown in Fig. 3B (top panel). This consistency indicates that the most informative network features (identified through feature selection), GCC node-degree variance and clustering coefficient, are robust and generalize well across repeated measurements. The robustness of these learned features is further validated by the model’s performance metrics, particularly the high mean F1 score (0.90) and its very low variance (0.001) across 10-fold cross-validation, as shown in Fig. 3B (bottom panel) highlighting both accuracy and stability in predictions.

## 4. DISCUSSION

Human brain organoids offer a powerful, scalable platform for modeling the complex cellular and network-level disturbances of genetic neurological disorders. While their potential is clear, the field has lacked comprehensive analytical pipelines that can translate the rich, dynamic data from these models into robust, interpretable biomarkers of disease. This work addresses that critical gap by introducing a comprehensive, time-resolved network analysis pipeline and demonstrating its efficacy in a tauopathy model. Our central finding is that dynamic graph-theoretic measures derived from two-photon calcium imaging can serve as a potent quantitative biomarker, reliably distinguishing *MAPT*-*mutant* assembloids from their isogenic *controls*. The success of our RF classifier, driven by only a few key network features, underscores the power of this approach. More importantly, the nature of these distinguishing features—specifically, the mean of the node degree variance and the clustering coefficient—provides deeper insight into the underlying pathophysiology. Our analysis revealed that *mutant* networks develop a topology characterized by the emergence of highly connected “hub” neurons and an increase in local interconnectedness. These are hallmarks of a network prone to hypersynchrony, a finding that parallels in vivo studies [10, 11] where impaired inhibitory function in models of Alzheimer’s Disease leads to epileptiform activity. The network topology we identified may, therefore, represent an in vitro correlate of a seizure-prone state, driven by the failure of inhibitory control that results in pathological synchronization. However, further studies are needed to specifically evaluate this.

The strength of this pipeline lies not only in its classificatory power but in its ability to generate testable, biologically relevant hypotheses. The observation of a “hub-like” network structure points toward specific cellular mechanisms that can now be explored, such as the potential vulnerability or dysfunction of inhibitory interneurons, which are known to be critical for regulating network synchrony. While this study is limited to a single *MAPT* variant and an in vitro system, our approach provides a versatile framework for studying a wide range of neurological disorders. Future investigation using this assembloid platform is necessary to define the cellular basis of this network dysfunction and to evaluate the specific role of interneuron pathology.

This assembloid-based pipeline offers several advantages for translational research over traditional rodent or 2D human models. By utilizing 3D patient-derived iPSC assembloids, we capture human-specific circuit organization and emergent dynamics without the need for non-physiologic gene overexpression. Furthermore, this platform provides a framework for precision medicine; iPSCs derived from individual patients can be used to generate “personalized” assembloids to evaluate specific network dysfunctions and test candidate therapeutic interventions in a patient-specific genomic background. This capacity for individualized functional profiling represents a significant step toward bridging the gap between preclinical discovery and clinical application.

## 5. CONCLUSION

We have developed and validated a novel pipeline for time-resolved network analysis of brain organoid models. Our approach successfully extracts a robust functional biomarker capable of distinguishing a pathogenic state from a healthy control with high accuracy. Crucially, the features that define this biomarker provide direct insight into the nature of the network dysfunction, bridging the gap between high-level dynamics and underlying cellular mechanisms. This work establishes a powerful methodology for dissecting the complex network alterations in human neurological diseases and for identifying quantitative targets for future cellular studies and therapeutic development.

## REFERENCES

[1] Giorgia Quadrato, Tuan Nguyen, Evan Z Macosko, John L Sherwood, Sung Min Yang, Daniel R Berger, Natalie Maria, Jorg Scholvin, Melissa Goldman, Justin P Kinney, et al., “Cell diversity and network dynamics in photosensitive human brain organoids,” Nature, vol. 545, no. 7652, pp. 48–53, 2017.

[2] Cleber A Trujillo, Richard Gao, Priscilla D Negraes, Jing Gu, Justin Buchanan, Sebastian Preissl, Allen Wang, Wei Wu, Gabriel G Haddad, Isaac A Chaim, et al., “Complex oscillatory waves emerging from cortical organoids model early human brain network development,” Cell stem cell, vol. 25, no. 4, pp. 558–569, 2019.

[3] Ranmal A Samarasinghe, Osvaldo A Miranda, Jessie E Buth, Simon Mitchell, Isabella Ferando, Momoko Watanabe, Thomas F Allison, Arinnae Kurdian, Namie N Fotion, Michael J Gandal, et al., “Identification of neural oscillations and epileptiform changes in human brain organoids,” Nature neuroscience, vol. 24, no. 10, pp. 1488–1500, 2021.

[4] Colin M McCrimmon, Daniel Toker, Marie Pahos, Qing Cao, Kevin Lozano, Jack J Lin, Jack M Parent, Andrew Tidball, Jie Zheng, László Molnár, et al., “Cortical versus hippocampal network dysfunction in a human brain assembloid model of epilepsy and intellectual disability,” Cell Reports, 2025.

[5] Madeline A Lancaster, Magdalena Renner, Carol-Anne Martin, Daniel Wenzel, Louise S Bicknell, Matthew E Hurles, Tessa Homfray, Josef M Penninger, Andrew P Jackson, and Juergen A Knoblich, “Cerebral organoids model human brain development and microcephaly,” Nature, vol. 501, no. 7467, pp. 373–379, 2013.

[6] Fikri Birey, Jimena Andersen, Christopher D Makinson, Saiful Islam, Wu Wei, Nina Huber, H Christina Fan, Kimberly R Cordes Metzler, Georgia Panagiotakos, Nicholas Thom, et al., “Assembly of functionally integrated human forebrain spheroids,” Nature, vol. 545, no. 7652, pp. 54–59, 2017.

[7] Shan Jiang, Natalie Wen, Zeran Li, Umber Dube, Jorge Del Aguila, John Budde, Rita Martinez, Simon Hsu, Maria V Fernandez, Nigel J Cairns, et al., “Integrative system biology analyses of crispr-edited ipsc-derived neurons and human brains reveal deficiencies of presynaptic signaling in ftld and psp,” Translational psychiatry, vol. 8, no. 1, pp. 265, 2018.

[8] Andrea Giovannucci, Johannes Friedrich, Pat Gunn, Jérémie Kalfon, Brandon L Brown, Sue Ann Koay, Jiannis Taxidis, Farzaneh Najafi, Jeffrey L Gauthier, Pengcheng Zhou, et al., “Caiman an open source tool for scalable calcium imaging data analysis,” elife, vol. 8, pp. e38173, 2019.

[9] M. E. J. Newman, Networks: An Introduction, Oxford University Press, Oxford, UK, 2010.

[10] Jorge J Palop, Jeannie Chin, Erik D Roberson, Jun Wang, Myo T Thwin, Nga Bien-Ly, Jong Yoo, Kaitlyn O Ho, Gui-Qiu Yu, Anatol Kreitzer, et al., “Aberrant excitatory neuronal activity and compensatory remodeling of inhibitory hippocampal circuits in mouse models of alzheimer’s disease,” Neuron, vol. 55, no. 5, pp. 697– 711, 2007.

[11] Laure Verret, Edward O Mann, Giao B Hang, Albert MI Barth, Inma Cobos, Kaitlyn Ho, Nino Devidze, Eliezer Masliah, Anatol C Kreitzer, Istvan Mody, et al., “Inhibitory interneuron deficit links altered network activity and cognitive dysfunction in alzheimer model,” Cell, vol. 149, no. 3, pp. 708–721, 2012.

